# Combination of antifungal drugs and protease inhibitors prevent *Candida albicans* biofilm formation and disrupt mature biofilms

**DOI:** 10.1101/2020.02.11.945030

**Authors:** Matthew B. Lohse, Megha Gulati, Charles S. Craik, Alexander D. Johnson, Clarissa J. Nobile

**Affiliations:** Department of Microbiology and Immunology, University of California – San Francisco, San Francisco, CA 94158; Department of Biology, BioSynesis, Inc., San Francisco, CA 94114; Department of Molecular and Cell Biology, University of California – Merced, Merced, CA 95343; Department of Pharmaceutical Chemistry, University of California – San Francisco, San Francisco, CA 94158

**Keywords:** *Candida albicans*, biofilms, antimicrobial resistance, therapeutics, protease inhibitors, aspartyl protease inhibitors

## Abstract

Biofilms formed by the fungal pathogen *Candida albicans* are resistant to many of the antifungal agents commonly used in the clinic. Previous reports suggest that protease inhibitors, specifically inhibitors of aspartyl proteases, could be effective antibiofilm agents. We screened three protease inhibitor libraries, containing a total of 80 compounds for the abilities to prevent *C. albicans* biofilm formation and to disrupt mature biofilms. The compounds were screened individually and in the presence of subinhibitory concentrations of the most commonly prescribed antifungal agents for *Candida* infections: fluconazole, amphotericin B, or caspofungin. Although few of the compounds affected biofilms on their own, seven aspartyl protease inhibitors inhibited biofilm formation when combined with amphotericin B or caspofungin. Furthermore, nine aspartyl protease inhibitors disrupted mature biofilms when combined with caspofungin. These results suggest that the combination of standard antifungal agents together with specific protease inhibitors may be useful in the prevention and treatment of *C. albicans* biofilm infections.

**Importance:** *Candida albicans* is one of the most common pathogens of humans. *C. albicans* forms biofilms, structured communities of cells several hundred microns thick, on both biotic and abiotic surfaces. These biofilms are typically resistant to antifungal drugs at the concentrations that are normally effective against free-floating cells, thus requiring treatment with higher drug concentrations that often have significant side effects. Here, we show that certain combinations of existing antifungal agents with protease inhibitors, including several drugs already commonly used to treat HIV patients, are effective at inhibiting biofilm formation by *C. albicans* and/or at disrupting mature *C. albicans* biofilms.

## Introduction

*Candida albicans* is a member of the human microbiota which asymptomatically colonizes the skin, mouth, and gastrointestinal tract of healthy humans (1–4). This fungal species is also one of the most common pathogens of humans, typically causing superficial dermal and mucosal infections (1, 5–11). When a host’s immune system is compromised (e.g. in patients undergoing chemotherapy or with AIDS), *C. albicans* can also cause disseminated bloodstream infections with mortality rates exceeding 40% (1, 12–15).

An important virulence trait of *C. albicans* is its ability to form biofilms, structured communities of cells several hundred microns thick, that can form on both biotic and abiotic surfaces (1, 4, 9, 16–19). When mature, these biofilms contain a mixture of yeast, pseduohyphal, and hyphal cells surrounded by an extracellular matrix (1, 3, 17–19). *C. albicans* forms biofilms on mucosal surfaces, epithelial cell linings, and on implanted medical devices, such as catheters, dentures, and heart valves (20, 21). Mature *C. albicans* biofilms also release yeast cells, which can seed new infections elsewhere in the host (22, 23).

*C. albicans* biofilms are typically resistant to antifungal drugs at the concentrations that are normally effective against planktonic (free-floating) cells, thus requiring higher drug concentrations, which can lead to host side effects, such as liver and kidney damage (20, 21, 24–27). Furthermore, *C. albicans* can also form polymicrobial biofilms with several companion bacterial species (28–35), further complicating treatment strategies. These polymicrobial biofilms can, for example, protect their bacterial inhabitants from environmental hazards (e.g. oxygen in the case of anaerobic bacteria) (36) and antibiotic treatments (e.g. protecting *Staphylococcus aureus* from vancomycin) (37–39). The drug-resistant nature of both single species and polymicrobial biofilms frequently makes removal of biofilm-infected medical devices the only treatment. However, this recourse is problematic when patients are critically ill or when device removal involves complicated surgical procedures (e.g. heart valve replacement) (20, 40, 41).

Currently, the three major classes of antifungal drugs used to treat *C. albicans* infections are the polyenes, azoles, and echinocandins (41, 42). The polyenes (e.g. amphotericin B) target ergosterol in the fungal cell membrane and are fungicidal against *C. albicans*. The azoles (e.g. fluconazole) inhibit the demethylase enzyme Erg11 from the ergosterol biosynthesis pathway and are fungistatic against *C. albicans*. Echinocandins (e.g. caspofungin), the most recently developed class of antifungal drugs, inhibit synthesis of the cell wall crosslinking component β-1,3-glucan and are fungicidal against *C. albicans*. Although novel derivatives within these classes have been introduced over the years, new classes of drugs have not been introduced. The limited size of the existing antifungals, both in terms of the distinct classes and in the number of drugs within several of these classes, creates several problems. As noted above, these classes of drugs typically have reduced effectiveness against biofilms relative to planktonic cells (20, 21, 24–27). Furthermore, long term exposure to these drugs, especially to members of the azole class, can give rise to antifungal resistance. Although the development of new antifungal agents is clearly called for, several recent *in vitro* studies have shown that combinations of antifungals with other extant drugs can be effective against *C. albicans* biofilms (43, 44).

Recently, we demonstrated the importance of several secreted proteases (Saps) for *C. albicans* biofilm formation (45, 46). Deletion of Sap5 and Sap6, both of whose expression is upregulated in biofilms (46), reduced biofilm formation *in vitro* and *in vivo* (45). Previous reports showed that treatment with aspartyl protease inhibitors, a class of drug commonly used to treat HIV patients, reduced the occurrence of oral candidiasis in immunocompromised patients independent of effects of the drug on the immune system through HIV remediation (47–49). Further work showed that several of the commonly used antiretroviral HIV aspartyl protease inhibitors could inhibit the Saps (50–57). Exposure to these protease inhibitors also reduced *C. albicans* adherence to materials commonly used in medical devices and to layers of host cells (58–60), although the magnitude of the latter effect differs greatly between distinct cell types (61). Aspartyl protease inhibitors have also been observed to reduce *C. albicans*-induced tissue damage, proliferation, and virulence *in vivo* in a rat vaginal model (54, 62). Finally, one study suggested that aspartyl protease inhibitors and the antifungal agents fluconazole or amphotericin B act synergistically against *C. albicans* in the planktonic form (63). To date, the studies of aspartyl protease inhibitors with regards to *C. albicans* emphasized their effects on planktonic cells. The one exception found that exposure to amprenavir, a common HIV antiretroviral protease inhibitor, could reduce *C. albicans* biofilm formation *in vitro* (57).

Given the number of protease inhibitors already approved for use in humans, including inhibitors of aspartyl proteases or other classes of proteases, we sought to evaluate the ability of a wide range of protease inhibitors to prevent (either alone or in combination with other antifungals) the formation of *C. albicans* biofilms or to act against mature biofilms. To evaluate the efficacy of these compounds in this regard, we screened three libraries containing 80 protease inhibitors in both biofilm inhibition and disruption assays. Each protease inhibitor was screened for biofilm efficacy individually and in combination with fluconazole, amphotericin B, or caspofungin. Although few of the protease inhibitors were effective against biofilms on their own, several, especially members of the aspartyl protease inhibitor class, were effective against biofilms when combined with either caspofungin or amphotericin B.

## Results

### Protease Inhibitor Libraries

We selected three libraries of protease inhibitors to screen for compounds with the abilities to inhibit and/or disrupt *C. albicans* biofilm formation *in vitro*. The first library, the SCREEN-WELL® Protease Inhibitor Library (Enzo Life Sciences), contains 53 protease inhibitors effective against several classes of proteases (File S1). The remaining two libraries contain 31 compounds known or predicted to specifically inhibit aspartyl proteases (64), of which we tested 27 in at least one assay. We focused on nine FDA-approved aspartyl protease inhibitors, developed to inhibit HIV-1 protease, ten macrocycles (API12-21), and eight linear peptidomimetics (API1-6, 10, and 11) that were originally synthesized with the goal of identifying new aspartyl protease inhibitors (64).

### Stand-alone Screens

We screened the three libraries for their abilities to inhibit biofilm formation or to disrupt mature biofilms using the Sustained Inhibition Biofilm Assay and Disruption Biofilm Assay (65, 66), respectively. In the Sustained Inhibition Biofilm Assay, compounds were included in media during the 90-minute adherence and 24-hour growth steps of the biofilm assay; the compounds were evaluated for their ability to reduce or prevent biofilm formation (Figure 1a). In the Disruption Biofilm Assay, a biofilm was grown for 24 hours before the compound of interest was added. The biofilm was then incubated for an additional 24 hours before determining whether the compound affected the mature biofilm (Figure 1a). In both assays, compounds were tested at a concentration of 40 μM.

**Figure 1:**
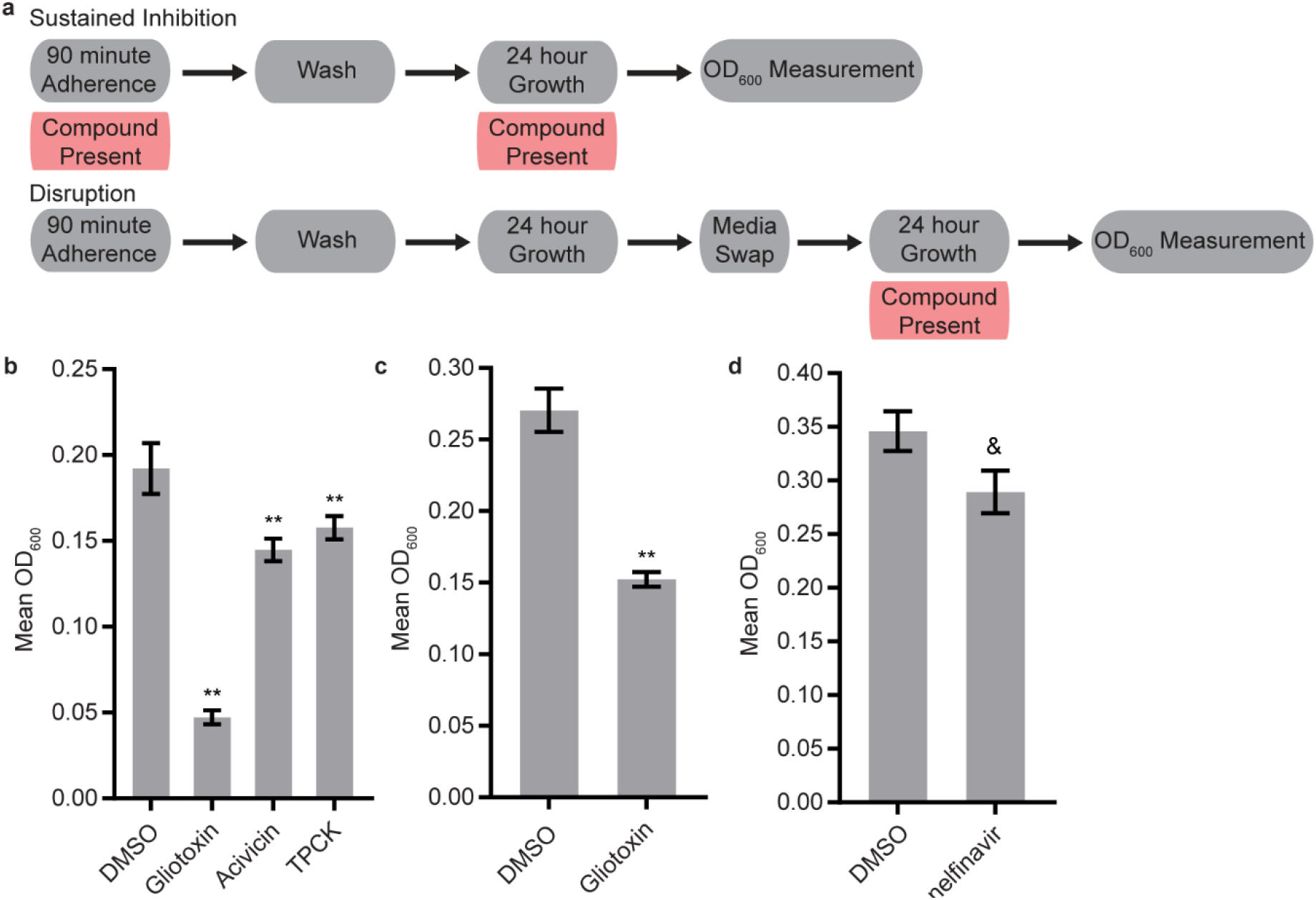
Four protease inhibitors either inhibited biofilm formation or disrupted mature biofilms on their own. (a) Overview of the experimental setups for the Sustained Inhibition and Disruption Biofilm Assays used for these experiments. For the Sustained Inhibition Biofilm Assay, compounds were included during both the 90-minute adherence step and the 24-hour growth step of a standard biofilm assay. For the Disruption Biofilm Assay, compounds were included during a second 24-hour growth step. (b-c) Statistically significant hits from the stand-alone (b) Sustained Inhibition and (c) Disruption assays with the SCREEN-WELL® Protease Inhibitor Library. Mean OD_600_ readings with standard deviations are shown; significant differences from the DMSO solvent control as determined by Welch’s t-test (two-tailed, assuming unequal variance) with the Bonferroni Correction are indicated for α=0.05 (*) and α=0.01 (**). Although a single repeat is shown, the indicated threshold was met by all of the repeats of each compound shown. (d) Statistically significant hit from the stand-alone Sustained Inhibition assays with the two aspartyl protease inhibitor libraries. Mean OD_600_ readings with standard deviations are shown; significant differences from the DMSO solvent control as determined by Welch’s t-test (two-tailed, assuming unequal variance) with the Bonferroni Correction are indicated. A single repeat is shown; the indicated significance threshold was met by two of the three repeats at α=0.01 while the third repeat did not pass at α=0.05. The “&” symbol indicates this mixed result.

Three of the 53 compounds in the SCREEN-WELL® Protease Inhibitor library, acivicin, gliotoxin, and TPCK, inhibited biofilm formation on their own (Figure 1b). One of these compounds, gliotoxin, also disrupted mature biofilms on its own (Figure 1c). TPCK irreversibly inhibits chymotrypsin (a serine peptidase) and can also inhibit some cysteine peptidases while gliotoxin inhibits the chymotrypsin-like activity of the 20S proteasome. Acivicin, on the other hand, is an inhibitor of gamma-glutamyl transpeptidase, an enzyme that transfers gamma-glutamyl groups from peptide donors to peptide acceptors as well as acting as a hydrolase to remove gamma-glutamyl groups from peptides. None of the 25 aspartyl protease inhibitors tested were able to disrupt mature *C. albicans* biofilms on their own, and only one of the 22 aspartyl protease inhibitors tested, the HIV-1 protease inhibitor nelfinavir, was able to inhibit biofilm formation on its own (BIC 50 μM) (Figure 1d).

### Combination Screens

We tested whether any compounds from the three protease inhibitor libraries could inhibit biofilm formation and/or disrupt mature biofilms in the presence of sub-inhibitory concentrations of amphotericin B, caspofungin, or fluconazole (see methods for concentrations). Five compounds from the SCREEN-WELL® Protease Inhibitor library inhibited biofilm formation in the Sustained Inhibition Biofilm Assay when combined with fluconazole (Figure 2a). We did not observe any synergies with amphotericin B or caspofungin in this assay. Two of these five compounds, gliotoxin and TPCK, were also “hits” in the stand-alone Sustained Inhibition Biofilm Assay described above. The remaining three compounds, lisinopril, Z-Prolyl-prolinal, and NNGH, were unique to the Sustained Inhibition Biofilm assay for synergies with fluconazole. Lisinopril inhibits the metalloprotease angiotensin-converting enzyme (ACE), NNGH inhibits matrix metalloproteinase 3 (MMP-3), and Z-Prolyl-prolinal inhibits prolyl endopeptidase (a serine protease). Two compounds from the SCREEN-WELL® Protease Inhibitor library, gliotoxin and Dec-RVKR-CMK, disrupted mature biofilms when combined with an antifungal agent (Figures 2b-c). Gliotoxin disrupted mature biofilms when combined with fluconazole (Figure 2c) while Dec-RVKR-CMK disrupted mature biofilms when combined with caspofungin (Figure 2b). Dec-RVKR-CMK, also known as furin convertase inhibitor, inhibits the subtilisin (Kex2p-like) proprotein convertase (a type of serine protease).

**Figure 2:**
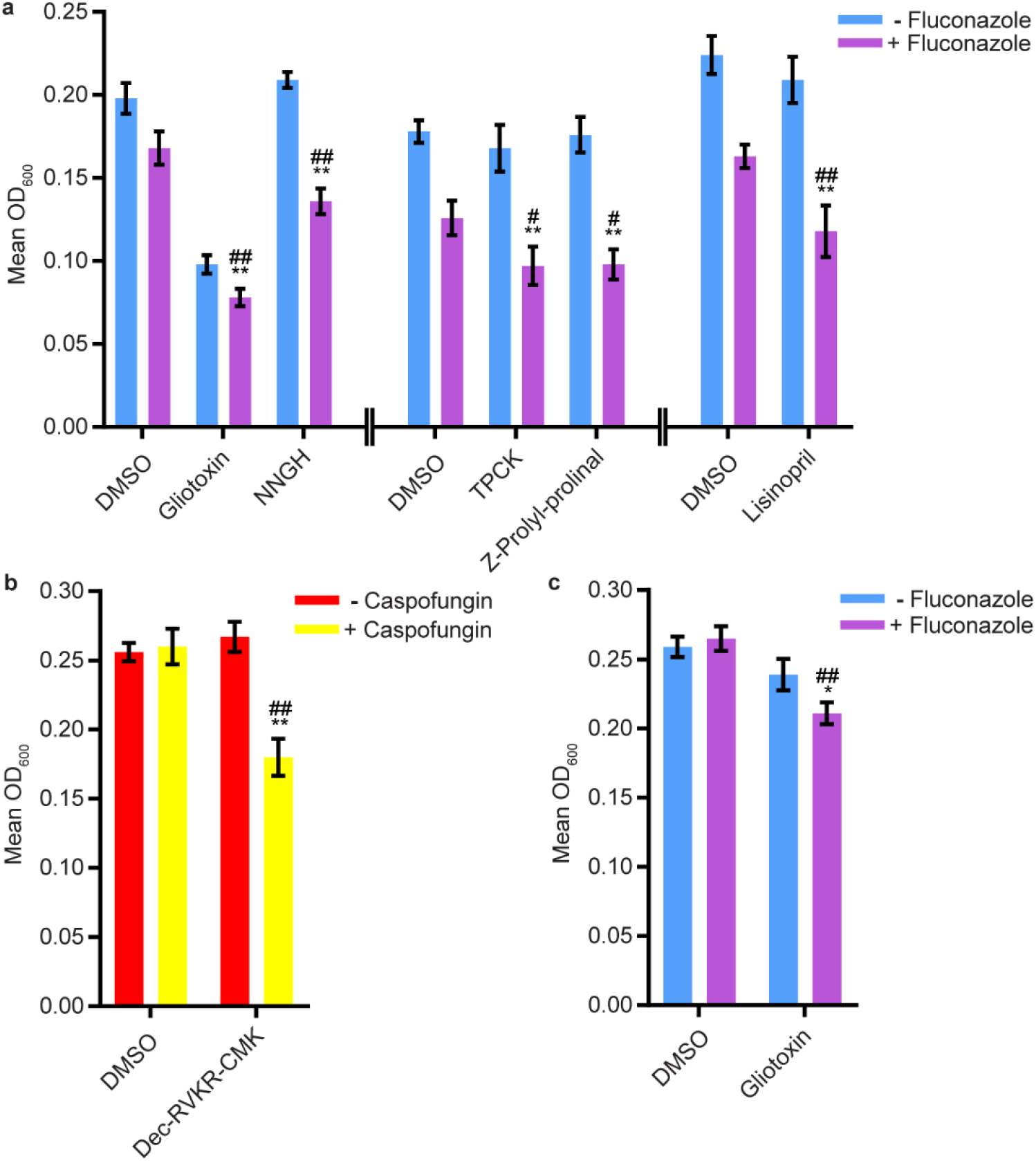
Six compounds from the SCREEN-WELL® Protease Inhibitor Library either inhibited biofilm formation or disrupted mature biofilms in combination with one or more antifungal agents. (a) Statistically significant hits from the combination Sustained Inhibition Biofilm Assays with fluconazole. For each compound, the wells with fluconazole (+ fluconazole) are indicated in purple and the wells without fluconazole (− fluconazole) are indicated in blue. (b) Statistically significant hits from the combination Disruption Biofilm Assays with caspofungin. For each compound, the wells with caspofungin (+ caspofungin) are indicated in yellow and the wells without caspofungin (− caspofungin) are indicated in red. (c) Statistically significant hits from the combination Disruption Biofilm Assays with fluconazole. For each compound, the wells with fluconazole (+ fluconazole) are indicated in purple and the wells without fluconazole (− fluconazole) are indicated in blue. For panels a-c, mean OD_600_ readings with standard deviations are shown; significant differences from the compound without antifungal agent control (e.g. gliotoxin, - fluconazole), as determined by Welch’s t-test (two-tailed, assuming unequal variance) with the Bonferroni Correction, are indicated for α=0.05 (*) and α=0.01 (**). Significant differences from the antifungal agent without compound control (e.g. DMSO, + fluconazole), as determined by Welch’s t-test (two-tailed, assuming unequal variance) with the Bonferroni Correction, are indicated for α=0.05 (#) and α=0.01 (##). Data from separate plates are separated by two vertical lines on the x-axis; the DMSO solvent control is shown for each plate.

We next evaluated 17 aspartyl protease inhibitors in the Sustained Inhibition Biofilm Assay and 26 aspartyl protease inhibitors in the Disruption Biofilm Assay in combination with the same three antifungal agents. Seven aspartyl protease inhibitors (four HIV-1 protease inhibitors and three macrocycles) inhibited biofilm formation when combined with one or more of the antifungal agents (six with caspofungin, five with amphotericin B, and one with fluconazole) (Figure 3). Specifically, lopinavir and API13 inhibited biofilm formation in combination with caspofungin while API19 inhibited biofilm formation in combination with amphotericin B. Ritonavir, saquinavir, and API15 inhibited biofilm formation in combination with caspofungin and amphotericin B while nelfinavir inhibited biofilm formation in combination with all three antifungal agents tested (Figure 3). Nine aspartyl protease inhibitors (the HIV-1 protease inhibitors atazanavir, indinavir, lopinavir, nelfinavir, ritonavir, saquinavir; and the macrocycles API15, API16, API19) disrupted mature biofilms in combination with caspofungin (Figure 4). None of the 26 aspartyl protease inhibitors tested disrupted biofilms in the presence of amphotericin B or fluconazole. We were surprised to find compounds that were effective at disrupting mature biofilms, but were not effective at inhibiting biofilm formation, namely atazanavir, indinavir, and API16. We also note that the macrocycle API19 had a synergistic effect with amphotericin B in the Sustained Inhibition Biofilm Assay but with caspofungin in the Disruption Biofilm Assay.

**Figure 3:**
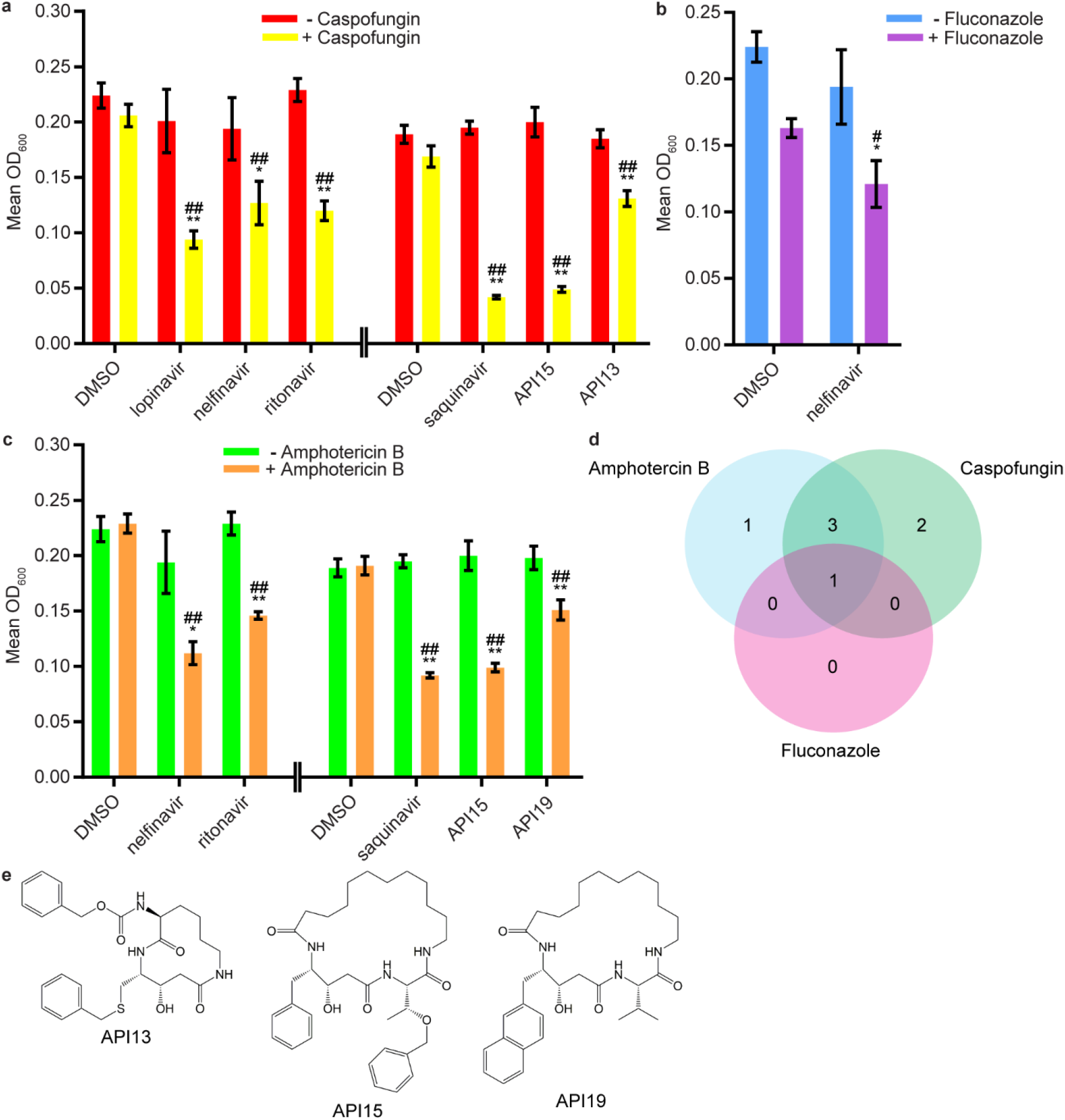
Seven aspartyl protease inhibitors were capable of inhibiting biofilm formation in combination with one or more of the three antifungal agents tested. (a) Statistically significant hits from the combination Sustained Inhibition Biofilm Assays with caspofungin. For each compound, the wells with caspofungin (+ caspofungin) are indicated in yellow and the wells without caspofungin (− caspofungin) are indicated in red. (b) Statistically significant hit from the combination Sustained Inhibition Biofilm Assays with fluconazole. For each compound, the wells with fluconazole (+ fluconazole) are indicated in purple and the wells without fluconazole (− fluconazole) are indicated in blue. (c) Statistically significant hits from the combination Sustained Inhibition Biofilm Assays with amphotericin B. For each compound, the wells with amphotericin B (+ amphotericin B) are indicated in orange and the wells without amphotericin B (− amphotericin B)are indicated in green. For panels a-c, mean OD_600_ readings with standard deviations are shown; significant differences from the compound without antifungal agent control (e.g. lopinavir, - caspofungin), as determined by Welch’s t-test (two-tailed, assuming unequal variance) with the Bonferroni Correction, are indicated for α=0.05 (*) and α=0.01 (**). Significant differences from the antifungal agent without compound control (e.g. DMSO, + caspofungin), as determined by Welch’s t-test (two-tailed, assuming unequal variance) with the Bonferroni Correction, are indicated for α=0.05 (#) and α=0.01 (##). Data from separate plates are separated by two vertical lines on the x-axis; the DMSO solvent control is shown for each plate. (d) Venn diagram illustrating the degree of overlap between the combination aspartyl protease inhibitor Sustained Inhibition Biofilm Assay screens with amphotericin B, caspofungin, or fluconazole. (e) Structure of the aspartyl protease inhibitors API13, API15, and API19.

**Figure 4:**
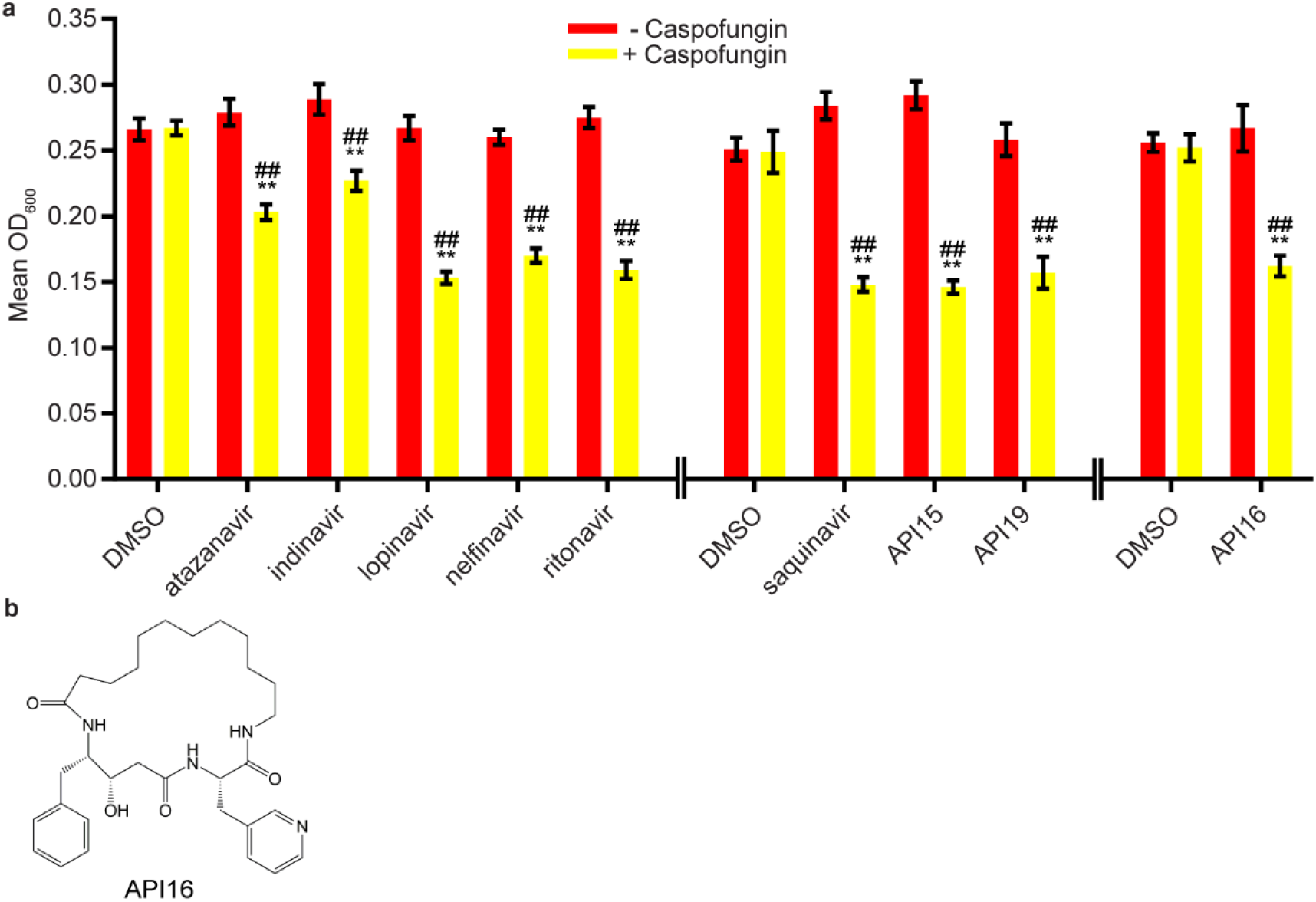
Nine aspartyl protease inhibitors disrupted mature biofilms in combination with the antifungal agent caspofungin. (a) Statistically significant hits from the combination Disruption Biofilm Assays with caspofungin. For each compound, the wells with caspofungin (+ caspofungin) are indicated in yellow and the wells without caspofungin (− caspofungin) are indicated in red. Mean OD_600_ readings with standard deviations are shown; significant differences from the compound without the caspofungin control (e.g. atazanavir, - caspofungin), as determined by Welch’s t-test (two-tailed, assuming unequal variance) with the Bonferroni Correction, are indicated for α=0.05 (*) and α=0.01 (**). Significant differences from the caspofungin without compound control (e.g. DMSO, + caspofungin), as determined by Welch’s t-test (two-tailed, assuming unequal variance) with the Bonferroni Correction, are indicated for α=0.05 (#) and α=0.01 (##). Data from separate plates are separated by two vertical lines on the x-axis; the DMSO solvent control is shown for each plate. (b) Structure of the aspartyl protease inhibitor API16.

## Discussion

The ability of *C. albicans* to form biofilms on biotic and abiotic surfaces presents a serious treatment challenge in the clinic as biofilms are typically resistant to all classes of antifungal drugs used to treat planktonic infections. Our results suggest that proteolysis is important for the maintenance of the *C. albicans* biofilm structure since anti-proteolytic agents contribute to the prevention and disruption of these biofilms. Proteases may play several different roles in *C. albicans* biofilm formation, an idea supported by the fact that proteases are dynamically expressed throughout the course of *C. albicans* biofilm formation (67, 68). For example, Sap5 and Sap6, two secreted aspartyl proteases that are highly upregulated at certain stages of biofilm formation, are known to mediate adhesion of *C. albicans* cells to surfaces (45) and possibly of *C. albicans* cells to one another. Proteases may also contribute to the breakdown and acquisition of nutrients, the processing of molecules important for biofilm formation (e.g. adhesion molecules), quorum sensing, and/or extracellular matrix production throughout biofilm formation and maintenance. Although the involvement of secreted proteases in biofilm formation is a relatively new concept, there is some precedent for this idea in bacterial biofilms, where extracellular proteases were found to be involved in the processing of adhesion molecules during biofilm formation of *Staphylococcus* species (69–71).

In this study, we identify several protease inhibitors from different classes that are effective at preventing biofilm formation and/or at disrupting established biofilms when combined with caspofungin, fluconazole, or amphotericin B, members of the three major antifungal classes used to treat fungal infections in the clinic. Aspartyl protease inhibitors, in particular those that inhibit HIV-1 protease, were the most effective compounds tested when combined with traditional antifungal agents. Combined with the known dependence on Sap5 and Sap6 for biofilm formation (45) and previous reports that aspartyl protease inhibitors affect *C. albicans in vitro* and *in vivo* (47–60, 62), aspartyl protease inhibitors are potentially promising combination treatments for *C. albicans* biofilm infections which are recalcitrant to single drug treatments. We note, however, that we screened fewer inhibitors of other classes of proteases than we did for aspartyl proteases. Despite this bias, we succeeded in identifying several inhibitors of two additional classes of proteases, serine and metalloproteases. It may prove rewarding to conduct additional screens of FDA-approved drugs whose mechanisms rely on the inhibition of other classes of proteases with the goal of repurposing these drugs as novel antifungals.

Perhaps the most unexpected result from this study was the identification of compounds capable of disrupting mature biofilms that were unable to prevent biofilm formation (Figure 5). Unlike the opposite case, where a compound that could prevent biofilm formation might be unable to penetrate a mature biofilm to have an effect, it is not readily apparent how the capacity to disrupt an established biofilm would not also inhibit the formation of a biofilm. Although we do not understand the basis for this result, it demonstrates that compounds that disrupt biofilms are not simply a subset of those that inhibit formation (Figure 5). This observation underscores the importance of screening compounds for their antibiofilm capabilities in both types of assays.

**Figure 5:**
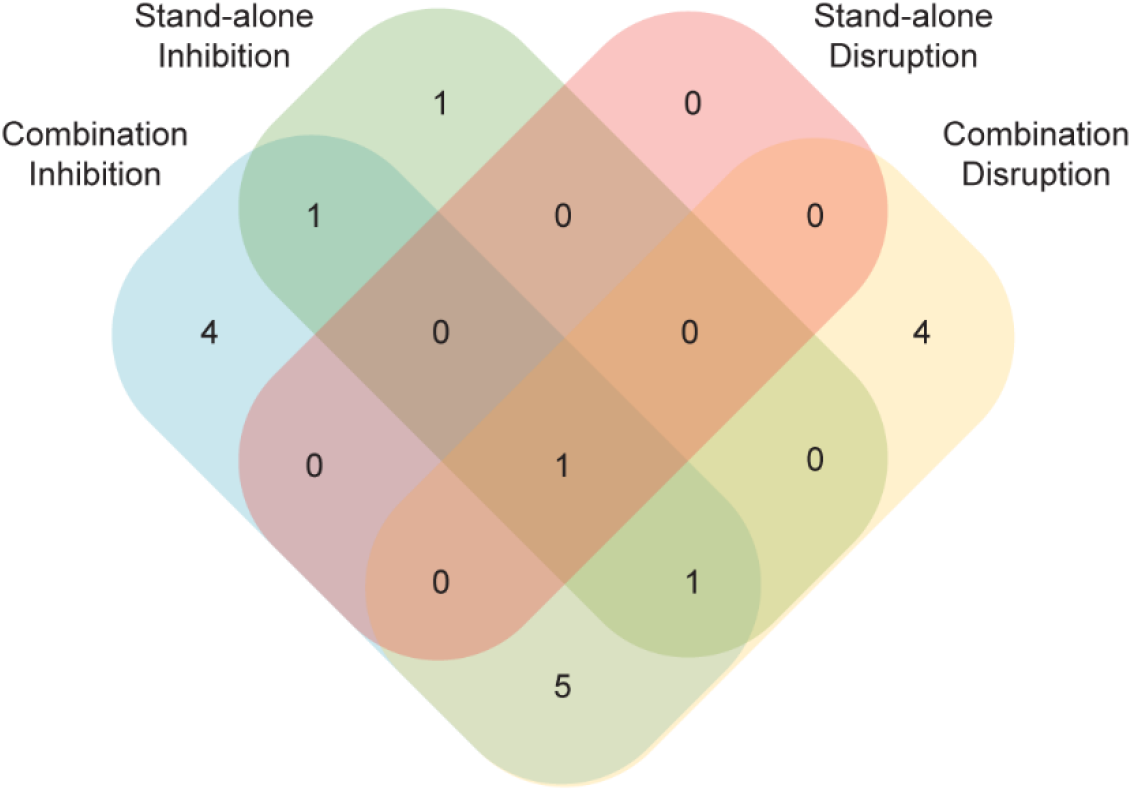
A number of compounds had effects in just a subset of the four biofilm assays. Compounds with an effect in either the stand-alone or the combination versions of the Sustained Inhibition or Disruption Biofilm Assays are indicated. In total, 17 compounds had an effect in at least one of the four assays.

Although we focused on one type of compound, protease inhibitors, this study raises several points to consider when screening for antibiofilm agents. First, consistent with previous reports (43, 44), our results highlight the importance of screening for synergistic interactions, as we detected more hits and hits with stronger effects against biofilms when existing antifungal agents were present along with the compound of interest (Figure 5). Second, our results highlight the importance of screening using biofilms as opposed to planktonic cultures. For example, in our biofilm assays with saquinavir, amphotericin B showed more synergy than fluconazole whereas the opposite relationship was reported for planktonic cultures (63). We also note that we identified compounds that had effects on their own but not in combination with existing antifungal agents, as well as the reverse. As such, pursuing multiple assays (e.g. planktonic versus biofilm, stand-alone compounds versus combinations) maximizes the chance of identifying useful compounds.

Finally, we note that this study was largely inspired by the discovery of the biofilm defects of the *sap5* and *sap6* single and double mutant strains (45). Thus, future compound library screening could be informed by other sets of gene knockouts with biofilm defects; likewise, results from chemical screens could identify genes (and their protein products) required for biofilm formation if the mechanism of action of the chemical compound is known. To further develop the idea of exploiting existing compounds, it should be possible to screen existing *C. albicans* mutant strain libraries for biofilm defects that arise in the presence of subinhibitory concentrations of traditional antifungal agents. Should biofilm formation by specific classes of mutant strains prove particularly sensitive to traditional antifungal agents, a subsequent combination screen between the traditional antifungal agents and compounds that affect that particular pathway of genes might prove informative.

## Materials and Methods

### Strains and Media

All assays were performed using SNY425, a SC5314-derived prototrophic a/α strain (72). *C. albicans* cells were cultured as previously described; in brief, cells were allowed to recover from glycerol stocks for two days at 30°C on yeast extract peptone dextrose (YEPD) plates (2% Bacto™ peptone, 2% dextrose, 1% yeast extract, 2% agar). Overnight cultures were grown approximately 16 hours at 30°C in YEPD media (2% Bacto™ peptone, 2% dextrose, 1% yeast extract). Biofilm assays were performed in RPMI-1640 media (containing L-glutamine and lacking sodium biocarbonate, MP Biomedicals #0910601) supplemented with 34.5 g/L MOPS (Sigma, M3183), adjusted to pH 7.0 with sodium hydroxide, and sterilized with a 0.22 μm filter (65, 66).

### Compound Libraries

The 53 member SCREEN-WELL® Protease Inhibitor Library (http://www.enzolifesciences.com/BML-2833/screen-well-protease-inhibitor-library/) was purchased from Enzo Life Sciences. The two aspartyl protease inhibitor libraries (from which we focused on nine FDA-approved HIV-1 protease inhibitors, the ten macrocycles, and eight linear peptidomimetics) have been previously reported (64). Due to limited quantities of several aspartyl protease inhibitors, a minority of compounds were only screened in one biofilm assay. In these cases, we prioritized the Disruption Biofilm Assay over the Sustained Inhibition Biofilm Assay. Four other compounds from these libraries (one FDA-approved HIV-1 protease inhibitor and three linear peptidomimetics (API7-9)) were not used in any assay. A list of compounds tested can be found in File S1.

### Biofilm Assays

The Sustained Inhibition and Disruption Standard Optical Density Biofilm Assays followed previously reported protocols for the 384-well format of biofilm screening assays (65–67, 73). Compounds and antifungal agents were added during the 90-minute adherence and 24-hour growth steps of the Sustained Inhibition Biofilm Assay or for the second 24-hour growth step of the Disruption Biofilm Assay. In brief, 1 μl of overnight culture was added to 90 μl media (or media with drug) in a well (final OD_600_ = 0.15, roughly 2×10^6^ cells/ml). Plates were then sealed with Breathe-Easy® sealing membranes (Diversified Biotech BEM-1) and shaken at 37°C for 90 min at 350 rpm in an ELMI (DTS-4) incubator. Media was removed, wells were washed with PBS, and fresh media (or media with drug) was added back to wells. Plates were then resealed and shaken for a further 24 hours. For the Sustained Inhibition Biofilm Assay, media was removed at this point and the absorbance (OD_600_) was determined on a Tecan Infinite M1000 Pro or a Tecan M200. For the Disruption Biofilm Assays, media was instead removed in groups of 6 to 12 wells and fresh media containing the compound of interest was carefully added back to the wells. Plates were then resealed and shaken for an additional 24 hours before removing media and recording absorbance as described above.

### Stand-alone Assays

Compounds were tested at 40 μM in both the Sustained Inhibition and Disruption Standard Optical Density Biofilm Assays (65, 66). Individual repeats of candidate compounds and DMSO solvent controls were performed in groups of eight wells. Each plate had between three and eight groups of control wells (24 to 64 total wells) spread throughout the plate to minimize position effects. For the SCREEN-WELL® Protease Inhibitor Library, the 53 compounds were screened once in both the Sustained Inhibition Biofilm Assay and the Disruption Biofilm Assay. Promising compounds from these initial screens were then tested a second time in the relevant assay(s). For the two aspartyl protease inhibitor libraries, we initially screened 21 compounds in the Sustained Inhibition Biofilm Assay and 25 compounds in the Disruption Biofilm Assay. Promising compounds from these initial screens were then tested two more times in the relevant assay(s). An additional three repeats were performed for four compounds (atazanavir, indinavir, nelfinavir, tipranavir) in the Disruption Biofilm Assay. For each experimental set of eight wells, significance was evaluated versus all of the control wells from the same plate by performing Welch’s t-test (two-tailed, assuming unequal variance). In order to correct for the multiple comparisons performed, we then applied the Bonferroni Correction with α = 0.05. All of the comparisons for a given type of assay (e.g. all of the stand-alone Sustained Inhibition Biofilm Assays) were pooled for this multiple comparisons correction step, giving a number of hypotheses, m, of 104 for the Sustained Inhibition Biofilm Assay and of 125 for the Disruption Biofilm Assay (final thresholds 4.81 × 10^−4^ and 4.00 × 10^−4^ respectively). We then determined whether each experimental repeat (1) had an average absorbance of less than the average of the control wells and (2) was significant after the multiple comparisons correction. To be considered a validated hit, a compound had to satisfy both of these criteria in each repeat if it was tested twice, for at least two repeats if it was tested three times, and for at least five repeats if it was tested six times. Data and statistics for the Stand-alone Sustained Inhibition and Disruption Optical Density Biofilm Assay are compiled in File S2. A summary of hits from these assays are included in File S3.

### BIC Assays

We determined the biofilm inhibitory concentration (BIC) of nelfinavir, tipranavir, and TPCK using the 384-well format Sustained Inhibition Standard Optical Density Biofilm Assay (65, 66). Both nelfinavir and tipranavir were serially diluted 2-fold from a maximum concentration of 200 μM to a minimum concentration of 0.1 μM. TPCK was serially diluted 2-fold from a maximum concentration of 512 μM to a minimum concentration of 0.06 μM. Groups of eight wells were used for each candidate compound or control condition and equivalent volumes of DMSO were used as loading controls for the compounds. Statistical testing was performed as described above with the following changes. Significance was evaluated for a given concentration of compound (e.g. 50 μM nelfinavir) compared to the equivalent DMSO loading control (e.g. the 50 μM loading control). All BIC comparisons were then pooled for multiple comparisons correction, giving a number of hypotheses, m, of 38 (α = 0.05, final threshold 1.32 × 10^−3^). We then determined whether each concentration of a drug (1) had an average absorbance of less than the average of the relevant control wells and (2) was significant after the multiple comparisons correction. The BIC of a compound was defined as the lowest concentration that met these requirements for which all higher concentrations of the same compound also met these requirements. If no concentration met these requirements, the BIC is indicated as greater than the highest concentration tested for that compound. Data and statistics for the BIC Sustained Inhibition Optical Density Biofilm Assay are compiled in File S2.

### Combination Assays

The combination (candidate compound plus known antifungal agent) Sustained Inhibition and Disruption Biofilm Assays followed the protocols described above with the following modifications. The candidate compounds were included at 12.5 μM in both assays with the exception of TPCK, Dec-RVKR-CMK, AEBSF·HCl, N-Ethylmaleimide, and acivicin, which were included at 4 μM, and gliotoxin, which was included at 1 μM. The Sustained Inhibition Biofilm Assays used 1 μg/mL amphotericin B, 0.125 μg/mL caspofungin, or 256 μg/mL fluconazole. The Disruption Biofilm Assays used 2 μg/mL amphotericin B, 0.5 μg/mL caspofungin, or 256 μg/mL fluconazole.

Compounds and controls were tested in groups of eight wells and two distinct groups of controls were included for all candidate compounds and antifungal agents tested on a given plate. The first set of controls contained the candidate compound, but no antifungal agent, while the second set of controls contained the antifungal agent, but no candidate compound. The concentration of candidate compound or antifungal agent in these control wells was the same as the experimental wells. In general, one set of wells was included for each experimental or control condition on a given plate. Statistical analysis was performed using Welch’s t-test and the Bonferroni Correction as described above with the following modifications. Each experimental condition was compared to both the relevant antifungal agent and candidate controls (e.g. a nelfinavir plus caspofungin experiment was compared to the nelfinavir-only control and the caspofungin-only control from the same plate). All of the same comparisons for a given assay (e.g. all of the Sustained Inhibition Biofilm Assay antifungal agent comparisons) were pooled for the multiple comparisons correction, giving a number of hypotheses, m, of 213 for both the antifungal agent and candidate comparisons in the Sustained Inhibition Biofilm Assay (α = 0.05, final threshold 2.35 × 10^−4^). The number of hypotheses, m, was 240 for both the antifungal agent and candidate comparisons in the Disruption Biofilm Assay (α = 0.05, final threshold 2.08 × 10^−4^). To be considered a hit, any given experimental condition must (1) have an average absorbance of less than the averages of both sets of relevant control wells and (2) remain significant for both sets of comparisons after the multiple comparisons correction. Data, statistics, and concentrations used for the combination Sustained Inhibition and Disruption Optical Density Biofilm Assay are compiled in File S2. A summary of hits from these assays are included in File S3.

## Acknowledgements

We thank Drs. Michael Winter and Starlynn Clarke for advice. This work was supported by National Institutes of Health (NIH) grants R43AI131710 (to M.B.L.), P50AI150476 (to C.S.C), R01AI083311 (to A.D.J.), and R35GM124594 and R41AI112038 (to C.J.N.). This work was also supported by the Kamangar family in the form of an endowed chair (to C.J.N). The content is the sole responsibility of the authors and does not represent the views of the funders. The funders had no role in the design of the study; in the collection, analyses, or interpretation of data; in the writing of the manuscript; and in the decision to publish the results.

## Author Contributions

Conceptualization, M.B.L., M.G., C.J.N. and A.D.J.; Methodology, M.B.L., M.G.; Validation, M.B.L.; Formal Analysis, M.B.L.; Investigation, M.B.L. and M.G.; Resources, C.S.C., C.J.N. and A.D.J.; Data Curation, M.B.L.; Writing – Original Draft Preparation, M.B.L.; Writing – Review & Editing, M.B.L., C.S.C., M.G., C.J.N. and A.D.J.; Visualization, M.B.L. and C.J.N.; Supervision, M.B.L., C.J.N. and A.D.J.; Project Administration, M.B.L., C.J.N. and A.D.J.; Funding Acquisition, M.B.L., C.S.C., C.J.N. and A.D.J.

## Supplemental File Captions

File S1: List of 80 compounds from the three protease inhibitor libraries tested in this study.

File S2: Compiled data and statistics from the Stand-alone and Combination Sustained Inhibition and Disruption Optical Density Biofilm Assays as well as the BIC Sustained Inhibition Optical Density Biofilm Assay. For each compound, the concentration used, average OD_600_, average OD_600_ of relevant control(s), and value(s) for Welch’s t-test versus the relevant control(s) are provided. Whether the average OD_600_ was below the average OD_600_ of the relevant control(s) and whether the difference from the relevant control(s) remains significant following the Bonferroni Correction (α=0.05) are also indicated.

File S3: Summary of the hits from the Stand-alone and Combination Sustained Inhibition and Disruption Optical Density Biofilm Assays.

